# Mammalian PIEZO channels rectify anionic currents

**DOI:** 10.1101/2024.08.23.609388

**Authors:** Tharaka D. Wijerathne, Aashish Bhatt, Wenjuan Jiang, Yun Lyna Luo, Jerome J. Lacroix

## Abstract

Under physiological conditions, mammalian PIEZO channels (PIEZO1 and PIEZO2) elicit transient currents mostly carried by monovalent and divalent cations. PIEZO1 is also known to permeate chloride ions, with a Cl^-^ / Na^+^ permeability ratio of about 0.2. Yet, little is known about how anions permeate PIEZO channels. Here, by separately measuring sodium and chloride currents using non-permanent counter-ions, we show that both PIEZO1 and PIEZO2 rectify chloride currents outwardly, favoring entry of chloride ions at voltages above their reversal potential, whereas little to no rectification was observed for sodium currents. Interestingly, chloride currents elicited by 9K, an anion-selective PIEZO1 mutant harboring multiple positive residues along intracellular pore fenestrations, also rectify but in the inward direction. Molecular dynamics simulation indicate that the inward rectification of chloride currents in 9K correlates with the largely positive electrostatic potential at the intracellular pore entrance, suggesting that rectification can be tuned by pore polarity. These results demonstrate that the pore of mammalian PIEZO channels inherently rectifies chloride currents.

**Statement of significance:** Mechanosensitive PIEZO ion channels play many important roles across cells and tissues. Their open pore facilitates the flow of cations down their electrochemical gradients, eliciting sodium-driven membrane depolarization and calcium-dependent signaling under physiological conditions. Yet, these channels also permeate chloride ions. In this study, we show that the two mammalian PIEZO channel homologs preferentially permeate chloride ions into the cells at voltages more positive than the chloride reversal potential. Although PIEZOs permeate cations more effectively than chloride ions, the influx of chloride ions mediated by PIEZOs could participate in certain physiological processes.

## Introduction

Mammalian PIEZO channels (PIEZO1 and PIEZO2) transduce mechanical stimuli into ionic fluxes that regulate important physiological functions across most cells, organs and tissues (1,2). Once opened, the pore of PIEZO1 channels facilitates the transmembrane diffusion of a wide range of cations down their electrochemical gradients. These include alkali ions (Na^+^ and K^+^), divalent ions (Ca^2+^ and Mg^2+^) as well as larger organic cations (3,4). PIEZO2 channels are thought to permeate a similarly large diversity of cations. For instance, recent evidence suggests the possibility that the open pore of PIEZO2 could permeate FM 1-43, a large fluorescent dye often used to label neurons (5). Besides cations, PIEZO1 has been shown to also permeate anions, with a Na^+^ vs. Cl^-^ permeability of about 0.2 (6).

We previously studied PIEZO1 permeation towards monovalent mineral ions *in silico* (7). This was done by measuring the rate of K^+^ and Cl^-^ ions crossing the pore of a computationally-generated open state under symmetric KCl solutions and under varying membrane potentials. In these conditions, our open state model generated a total conductance and monovalent cations vs. anions selectivity similar to experimental values. In addition, while potassium ions permeated equally well in both directions, chloride ions permeated more effectively in the inward direction (i.e., at voltages above the 0 mV reversal potential) than in the outward direction (i.e., at negative voltages), suggesting that PIEZO1 rectifies chloride, but not potassium currents. This result was surprising because the total PIEZO currents measured in physiological saline is not known to exhibit significant rectification (8).

Here, to test these computational predictions experimentally, we separately measure sodium and chloride currents using excised patches and whole-cell electrophysiology in presence of symmetrical solutions containing non-permeant counterions. Our results show that both wild-type PIEZO1 and PIEZO2 channels mediates outwardly-rectifying chloride currents whereas their sodium currents display little to no rectification, as predicted from our simulations. In addition, our experiments and simulations show that 9K, an anion-selective PIEZO1 mutant in which 6 neutral (S2491, S2150, N2151, C2154, I2164, S2168) and 3 acidic (E2172, E2491, E2496) intracellular residues along intracellular pore fenestrations are replaced by lysine (6), also rectifies chloride currents but in the inward direction (6). These results show that our computational PIEZO1 open state model faithfully recapitulates PIEZO1’s permeation properties, demonstrates that mammalian PIEZO channels inherently rectify chloride currents, and suggest that ionic rectification can be tuned by the electrostatic potential along the channel’s permeation pathway.

## Materials and Methods

### Patch-clamp electrophysiology

Plasmids encoding WT mouse PIEZO1 or the 9K PIEZO1 mutant were generously gifted by Ardem Patapoutian (Scripps Research, La Jolla) and Bailong Xiao (Tsinghua University, Beijing), respectively. These plasmids were used to transfect fibroblastic HEK293T cells as previously described (9-11). Cells were re-seeded on glass coverslips coated with Matrigel (Corning) one day after transfection and used for experiments one day after. Patch pipettes were pulled from G150F borosilicate capillaries (Warner Instruments) to a resistance of 2-3 MΩ using a P-97 puller (Sutter Instrument) and heat-polished using a Microforge-MF2 (Narishige). Patch pipettes were filled with 10 mM EGTA, 10 mM HEPES and 150 mM of either N-methyl-D-glucosamine chloride (NMDG-Cl) or sodium D-Gluconate (Na-GLU) (all chemicals were purchased from MilliporeSigma). The pH of both solutions was adjusted to 7.4 using NMDG/HCl (for the NMDG/Cl solution) and Gluconic acid/NaOH (for the Na-GLU solution). Osmolarities of these solutions was measured at 290-310 mOsm L^-1^.

Inside-out patches were formed by rapidly pulling the patch pipette away from the cell after gigaohm seal formation. Pressure pulses were delivered to excised-patches through the patch pipette using a high-speed pressure clamp (ALA Scientific). Pressure-induced currents were recorded at 10 kHz in the voltage-clamp mode using an Axopatch 200B amplifier and digitized using a Digidata 1550B (Molecular Devices). After confirming the presence of pressure-induced currents, HBSS bath solution (facing the intracellular side of the membrane) was replaced by perfusing NMDG-Cl or Na-GLU onto the patch using an 18-gauge blunt canula attached to an 8-channel pressurized perfusion system (AM-PS8-PR, Sutter Instrument). 20 seconds after the onset of perfusion, pressure-activated (−80 mmHg) currents were measured at +90, +70, +50, +30, +10, 0, -10, -30, -50, -70 and -90 mV in the presence of continuous perfusion. Patches were held at +5 mmHg and 0 mV for 20 s between each measurement to allow complete recovery from inactivation (9,12). An Ag/AgCl (3M KCl) reference electrode was used as bath ground. Junction potentials, measured in excised patches from untransfected cells in the current-clamp mode, were systematically below 2 mV and thus were not corrected from I-V curves.

Poking stimuli were provided to whole cells using a blunt glass probe attached to a piezoelectric actuator (P-841, Physik Instrumente) controlled by Clampex via an amplifier (E-625, Physik Instrumente). PIEZO2 currents were activated by a 3 μm poking stimulus. Poking distance was calibrated before each experiment as described before (9,11,13). After allowing ∼1 min for dialysis of intracellular fluid with patch pipette saline, currents were measured at +90, +70, +50, +30, +10, 0, -10, -30, -50, -70 and -90 mV. Patches were held at 0 mV for 20 s between each measurement to allow complete recovery from inactivation (12).

### Atomistic simulations

The PIEZO1 pseudo 9K mutant was prepared using our previous PIEZO1 open state model obtained from Anton2 simulation (7). Instead of changing the residue types, we altered the charge of the side chains of S2150, N2151, C2154, I2164, S2168, E2172, S2491, E2495, and E2496 (SNCISESEE) to mimic the charge of 9K mutant. This pseudo 9K mutant allowed us to investigate the effect of protein electrostatics on ion permeation without altering any covalent bonds and vdW interactions. To maintain the system neutral, we added an extra 36 Cl^-^ ions in the 150 mM KCl bulk solution. Only the central pore domain and repeat A (residues 1976–2546) were used for conductance simulation. The total system contains 622,323 atoms, including 160539 water molecules and 826 POPC lipid molecules. The voltage simulation was conducted using the GROMACS (version 2023.3) using the Charmm36m force field. The van der Waals interactions were truncated using a cutoff value of 1.2 nm and the force-switching function was applied between the distance range 1.0-1.2 nm. Long-range electrostatics were evaluated using the particle mesh Ewald method and a 1.2 nm cutoff was employed for short range electrostatics. All bonds containing hydrogen atoms were constrained using the LINCS algorithm. To prevent conformational change induced by artificial charged residues, positional restraints of 500 kJ/mol/nm2 was applied to the protein backbone during simulation. The system underwent minimization and equilibration in constant volume and temperature (NVT) and then in constant pressure and temperature (NPT) condition. The Nose-Hoover thermostat and Parrinello-Rahman barostat were employed to maintain 310.15 K and 1 atm during the production run. To compute the current-voltage curves for K+ and Cl-, constant external electric fields were applied in the z-direction. Two replicas of 100 ns simulation was run at each voltage (−0.5 V, -0.25 V, 0.25 V, and 0.5 V). Ionic current was measured by counting the total number of ions flowing through the pore from the extra cellular to the cytoplasmic side and vice versa. Visualization of ionic density in the pore was done using the VMD volmap plugin using an isosurface density cutoff of ± 0.009 e/Å3. The electrostatic potential was computed using PBEQ-Solver in CHARMM-GUI (14). Default dielectric constants (80 for solvent, 1 for protein) and salt concentration of 0.15 M were used.

## Results

To measure PIEZO currents specifically carried by sodium or chloride ions, we sought to use salts containing large, non-permeant counter-ions. We chose sodium gluconate (Na-GLU) to measure sodium currents and N-methyl-D-glucosamine chloride (NMDG-Cl) to measure chloride currents. We first tested whether GLU and NMDG could produce any detectable currents through wild-type PIEZO1 and through the anion-selective 9K variant. To this aim, we excised inside-out membrane patches (from untransfected and transfected HEK293T^PIEZO1KO^ cells) using patch pipettes filled with 150 mM NMDG-GLU. To verify that our patches contain functional channels, we first measured currents evoked by a -80 mmHg suction pulse in presence of a 150 mM NaCl bath solution. Patches excised from cells transfected with wild-type (WT) PIEZO1 tend to produce large outward currents at positive potentials, as expected if these currents are mainly carried by sodium ions. By contrast, patches excised from cells transfected with 9K produce large inward currents at negative potentials, as expected if these currents are mainly carried by chloride ions (**Fig. 1a**). These results demonstrate that suction-evoked currents from our excised patches are specifically mediated by PIEZO1 and its 9K variant.

**Figure 1.**
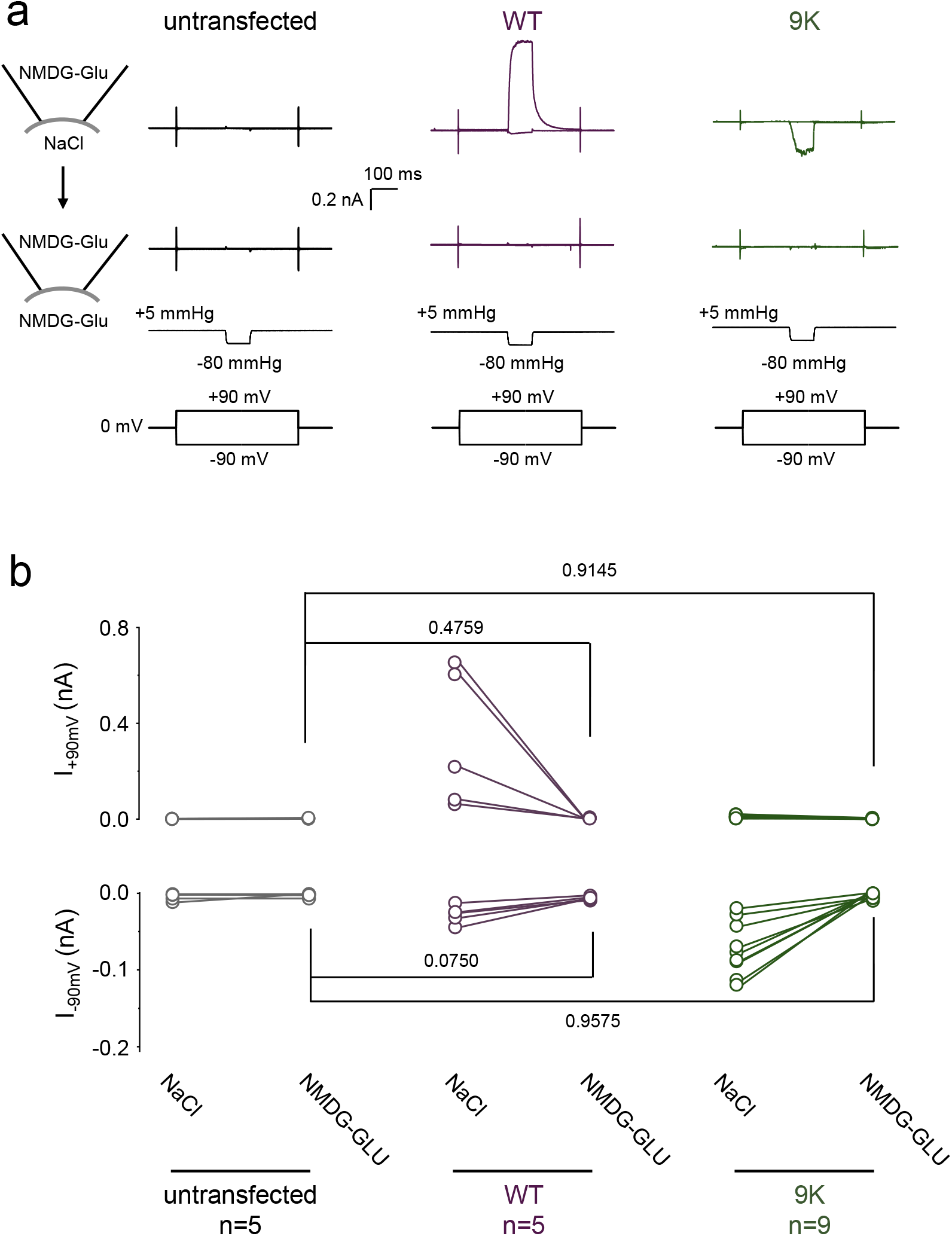
Neither gluconate nor NMDG permeates through PIEZO1 or the 9K mutant. (**a**) Representative stretch-activated currents traces measured from untransfected (left), WT PIEZO1 (middle) and the 9K mutant (right) membrane-excised patches before (top) and after (bottom) replacing the 150 mM NaCl bath (internal) solution with 150 mM NMDG-GLU. (**b**) Currents recorded from untransfected cells (left), or cells transfected with WT PIEZO1 (middle) or the 9K mutant (right) before and after replacing the bath solution. The numbers above plots are exact p-values from Kruskal-Wallis with Dunn’s multiple comparison tests.

After confirming the presence of these PIEZO-specific currents, we subsequently replaced the bath (internal) solution with 150 mM NMDG-GLU and repeated our recording protocol. Removing NaCl from the bath completely eliminates all currents in all tested patches, showing that neither GLU nor NMDG permeates through the open pore of WT PIEZO1 or 9K (**Fig. 1b**).

Since neither GLU nor NMDG permeates the channel pore, we repeated our measurements in the presence of symmetrical Na-GLU or NMDG-Cl to study the permeation of Na^+^ and Cl^-^ ions in both WT PIEZO1 and 9K (**Fig. 2a-d**). In WT PIEZO1, sodium currents tend to inactivate more rapidly at negative voltages, which is consistent with the known effect of voltage on PIEZOs’ inactivation kinetics (8). The peak amplitude of sodium currents follows a nearly linear relationship with respect to the applied voltage, but the peak amplitude of chloride currents did not: although chloride currents are clearly detectable at voltages above the reversal potential, they nearly vanish at voltages below it. Conversely, the 9K mutant displays almost no detectable sodium current and its chloride currents display robust inward rectification.

**Figure 2.**
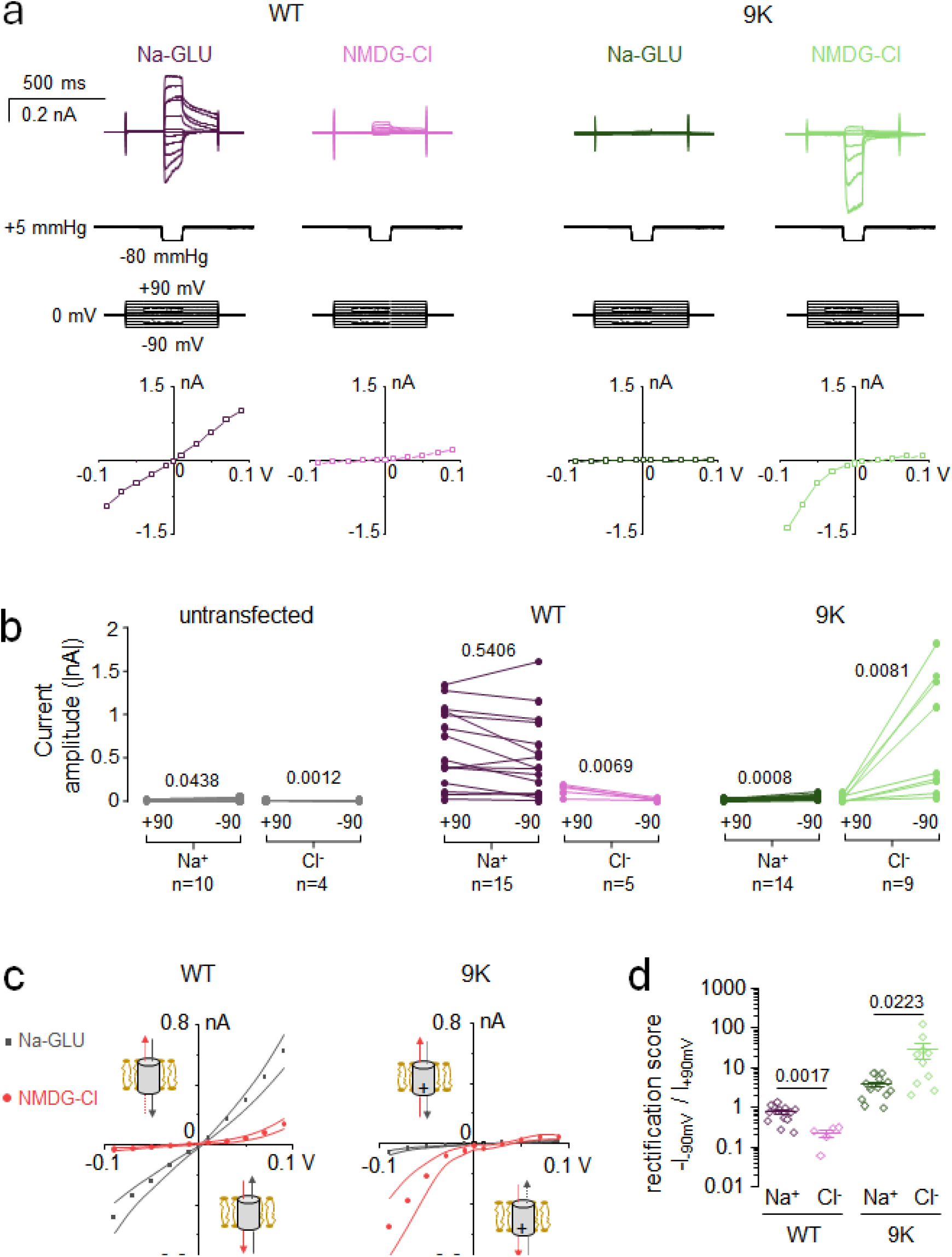
Rectification of chloride currents in PIEZO1 channels. (**a**) Exemplar inside-out excised patch recordings and corresponding I-V plots for WT PIEZO1 and the 9K mutant in the presence of symmetrical Na-GLU or NMDG-GLU solutions. (**b**) Comparison of current amplitude measured at ±90 mV for indicated experimental conditions. (**c**) I-V curves for WT and 9K measured in the presence of Na-GLU (grey) or NMDG-Cl (red) from data shown in (b). Lines represent s.e.m. (**d**) Rectification scores plotted for each excised patch for sodium and potassium currents in both WT and 9K. Numbers above plots in panels (b) and (d) indicate exact p-values from paired and two-tail T-tests, respectively.

A pairwise comparison of peak current amplitudes at -90 mV vs. +90 mV shows that, in WT PIEZO1, the amplitude of sodium currents is independent of the voltage polarity (p-value = 0.5406). However, the amplitude of chloride currents tend to decrease to nearly zero at negative voltages (p-value = 0.0069) (**Fig. 2b**). In the anion-selective 9K mutant, the amplitude of sodium currents is too low to determine whether they rectify. However, the amplitude of chloride currents is much larger at -90 vs. +90 mV (p-value = 0.0081). These results are recapitulated in current vs. voltage (I-V) curves (**Fig. 2c**). Rectification scores were calculated by dividing the absolute current amplitude at -90 mV by the current amplitude measured at +90 mV (**Fig. 2d**). In WT PIEZO1, the rectification score for sodium currents is near unity (0.78 ± 0.08), whereas it decreases to 0.23 ± 0.05 for chloride currents (p-value = 0.0017), indicating outward rectification. In the 9K mutant, the rectification score for chloride currents is 28.87 ± 12.20, revealing the extreme rectification behavior of the mutant. Sodium currents seem to rectify in the inward direction as well, although it seems difficult to interpret these results since the amplitude of sodium currents remains very low even at negative voltages. Taken together, our results overall confirm our computational predictions (7).

We next sought to test if the rectification phenotype of the 9K mutant is directly caused by the large excess of positive charges along intracellular pore fenestrations, which amounts to +36*e* for the trimeric channel compared to WT PIEZO1. To this aim, we computationally created a pseudo 9K mutant by appending a +1*e* charge to each mutated residue using our computational open state model, and evaluated ionic permeation properties under symmetrical KCl. The computational I-V curves recapitulate the main features of experimental I-V curves for the 9K mutant: robust chloride currents that rectify inwardly and faint sodium currents that are only detectable at negative voltages (**Fig. 3a**, for comparison, the I-V curves for WT PIEZO1 generated from our previous study is also plotted). The intracellular fenestrations of the pseudo 9K mutant shows a highly positive electrostatic potential compared to WT PIEZO1 (**Fig. 3b**, red circle). As expected, a larger density of chloride ions gather around the intracellular fenestrations in the pseudo 9K mutant compared to WT PIEZO1, likely due to the excess intracellular positive potential. The large excess of positive charges thus promotes the interaction of chloride ions with the intracellular pore fenestrations (**Fig. 3c**). This might favor the outward flow of chloride ions at negative voltages, consistent with the inward rectification of the 9K mutant. In addition, the pore of the pseudo 9K mutant displays a lower density for potassium ions compared to WT PIEZO1 (**Fig. 3c**), consistent with the very low amplitude of cation currents displayed by the 9K channel (**Fig. 2**).

**Figure 3.**
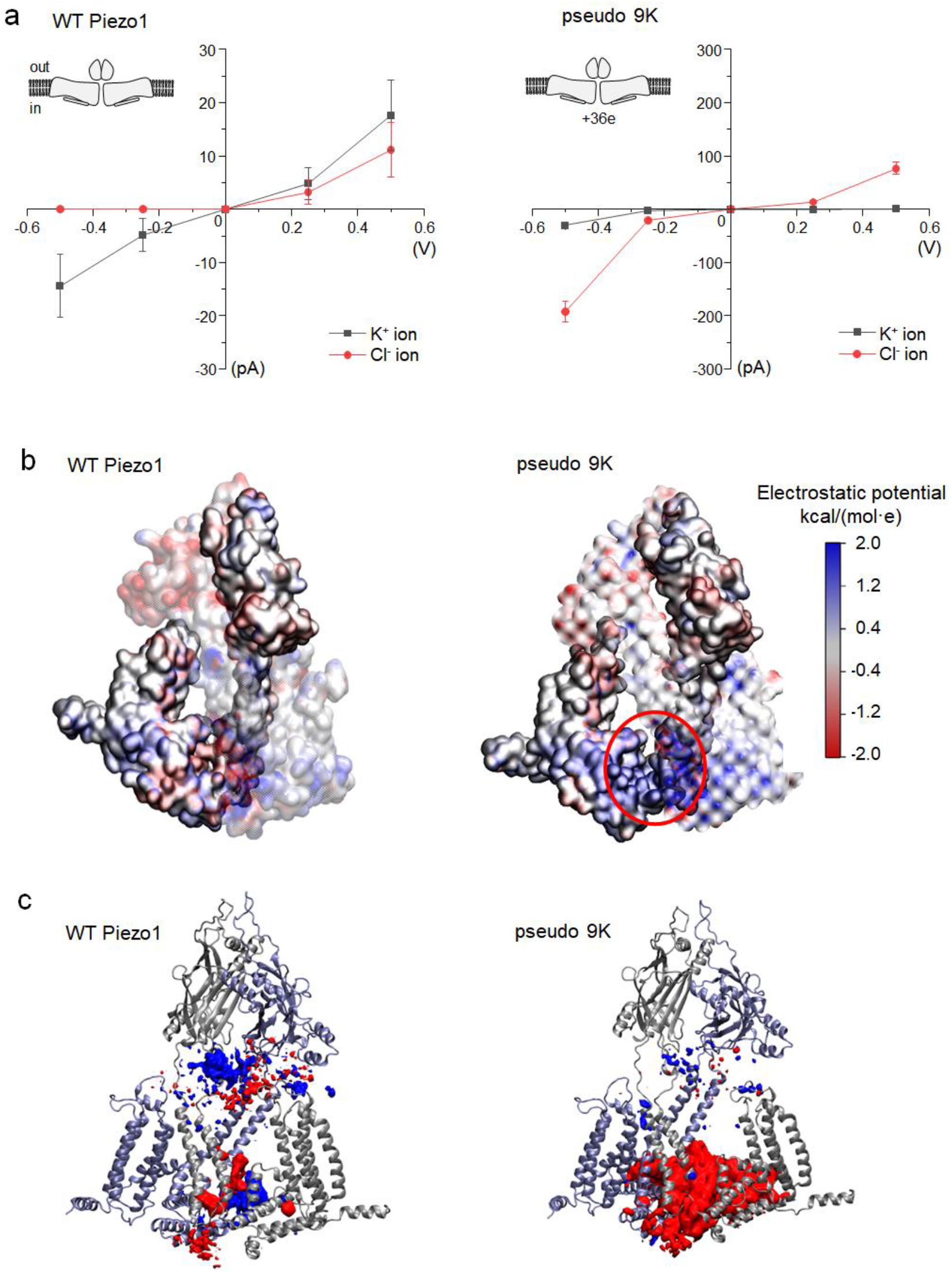
Computational electrophysiology predicts rectification and ionic selectivity in PIEZO1 WT and in the 9K mutant. (**a**) *Left*: I-V curves for wild-type PIEZO1 channel, reproduced from Jiang et al., 2021 (7). *Right*: I-V curves for the pseudo 9K mutant. Error bars are standard deviations from two independent replicas. (**b**) Electrostatic potential surface of WT and pseudo 9K mutant. Only two pore subunits are shown (in surface mode) for clarity. 9K residues are located in the red circled area. (**c**) Densities of potassium (blue) and chloride (red) ions within 18 Å of the pore in the wild-type and 9K systems averaged over 25 ns (250 frame). Two pore subunits are shown (in Newcartoon mode) in silver and iceblue.

We finally sought to test if the rectification behavior observed in PIEZO1 also extends to its homolog PIEZO2. We initially tried measuring PIEZO2 currents from excised patches but the current amplitude was too low to enable meaningful interpretations. We instead used the whole-cell poking technique to measure PIEZO2 currents, allowing the interior of the cell to be dialyzed by the pipette solution after breaking-in. To test if internal cations have been effectively dialyzed, we first tested the presence of PIEZO2 currents by poking cells in the presence of NMDG-GLU in the pipette and NaCl in the bath at -90 mV and +90 mV (**Fig. 4a-b**). In these conditions, we only detected inward currents consistent with entry of sodium ions at -90 mV. These inward currents were completely eliminated by subsequently replacing the bath solution with NMDG-GLU, showing that internal cations have been effectively dialyzed.

**Figure 4.**
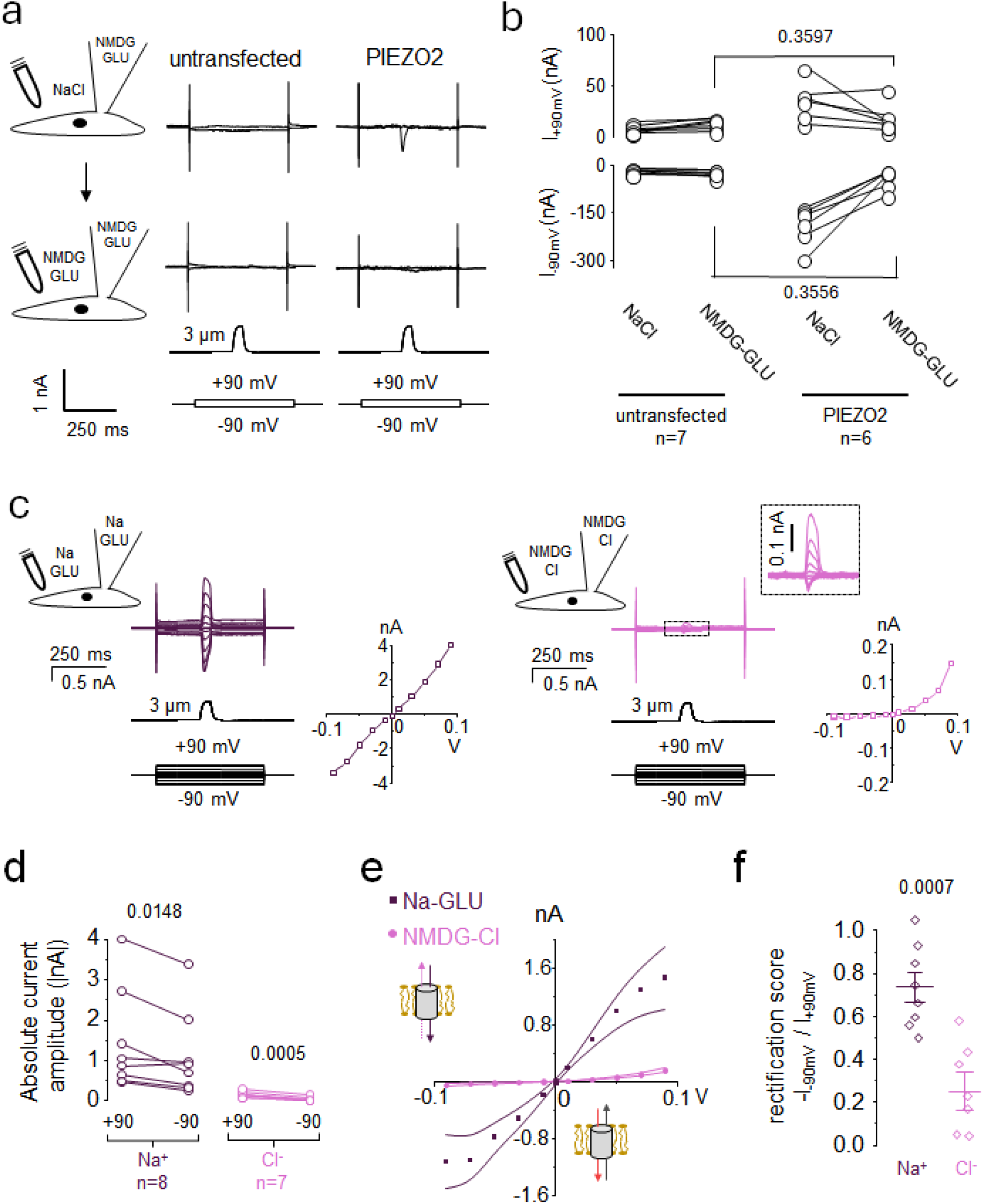
Chloride rectification in PIEZO2 channels. (**a**) Representative poking-induced whole-cell current traces measured at + 90 and -90 mV from untransfected and WT PIEZO2 cells before (top) and after (bottom) replacing the bath NaCl solution with NMDG-GLU. (**b**) Pairwise comparison of peak current amplitude from experiments shown in (a). (**c**) Exemplar whole cell recordings and corresponding I-V plots for PIEZO2 in the presence of symmetrical Na-GLU or NMDG-GLU solutions. (**d**) Comparison of current amplitude measured at ±90 mV from experiments shown in (c). (**e**) I-V curves for PIEZO2 in the presence of Na-GLU (grey) or NMDG-Cl (red) from data shown in (c). Lines represent s.e.m. (**f**) Rectification scores plotted for each cell for sodium and potassium currents. The numbers above plots indicate p-values from either paired two-tail T-tests(panels b and d) or two-tail T-test (panel f).

In the presence of symmetric Na-GLU, PIEZO2 elicits poking-dependent sodium currents whose peak amplitude appear more or less linear to the electromotive force, exhibiting a rectification score near unity (**Fig. 4c-f**). But in the presence of symmetrical NMDG-Cl, the peak chloride currents elicited by PIEZO2 in response to a poke stimulus clearly rectify in the outward direction. Hence, the rectification properties of PIEZO2 towards sodium and chloride currents are overall similar to that of PIEZO1.

## Discussion

Rectification is a general electrodynamic phenomenon in which the voltage-driven displacement of ions through confined nanopores is favored in one direction over the other. Many biological nanopores formed by channel proteins embedded into cell membranes exhibit rectification. Known biological mechanisms of ion channel rectification include voltage-dependent pore block by non-permeant solutes (15) and voltage-dependent conformational changes that alter the ability of the pore to permeate ions (16).

Yet, besides these well-known rectification mechanisms, many evolutionary-unrelated ion channels rectify currents with no evidence of conformational changes or pore-block (17-21), suggesting that rectification can also result from inherent biophysical properties of the channel pore. This study suggests that this may also be the case for mammalian PIEZO channels. Indeed, PIEZOs’ ability to rectify chloride currents outwardly is unlikely to result from a voltage-dependent pore blocking mechanism or substantial change of open probability, as such mechanisms should have caused sodium currents to rectify in a similar fashion. Yet, we did observe that, for both PIEZO1 and PIEZO2, sodium currents marginally rectify in the outward direction, their amplitude being slightly smaller at -90 mV compared to +90 mV. This apparent rectification behavior could be caused by PIEZOs’ faster inactivation kinetics at negative voltages (8,22): at these voltages, the faster inactivation would cause PIEZO currents to decay earlier, reducing the current peak amplitude.

The physical basis of intrinsic rectification has been investigated in the bacterial toxin alpha-hemolysin (23-26) and in synthetic nanopores made from organic and inorganic polymers (27-31). These studies broadly support the idea that rectification can be caused by the presence of asymmetric charges along the ion conduction pathway. Our study shows that the pore of mammalian PIEZOs rectifies chloride currents while sodium currents exhibit little to no rectification, suggesting that rectification can discriminate among permeant ions based on specific pore-ion electrostatic interactions. Our simulations support an electrostatic mechanism in which a large density of positive charges promotes the binding of chloride ions, which may facilitate their outward vs. inward flow when measured at driving forces of opposite signs but equal amplitude.

Our simulations results recapitulated the near linear cationic conductance and outward-rectifying anionic conductance seen in experiments. Yet, the relative amplitude of anionic vs. cationic currents at positive voltages, shown in **Fig. 3a**, was notably larger than experimentally observed in this study and others (6). This discrepancy could be due to the fact that our simulations used K^+^ ions as cations instead of Na^+^ ions, and/or to the fact that both K^+^ and Na^+^ ions permeate simultaneously in our simulations whereas they permeate separately in our experiments, and/or to the larger membrane potentials applied during simulations to speed up conductance.

It is difficult to determine whether the ability of PIEZO channels to rectify chloride currents confers a specific evolutionary advantage, or if this property has emerged without a particular biological role. One possible advantage could be the reduction of unwanted chloride currents at more negative voltages. This scenario is likely to be met at the physiological resting potential of excitable cells, which tends to lie either near or below the chloride reversal potential (32-34). Another possible role could be to promote the entry of chloride ions at more positive potentials. Although we were unfortunately not able to detect PIEZO1 single channel chloride currents in excised patches, PIEZO1 unitary conductance for chloride ions is expected to be about 15-20% of the sodium conductance (∼68 pS in absence of calcium ions) (6), which would be within or above the range of chloride conductance exhibited by other chloride selective pores (1-10 pS) such as ClC channels and CFTR (35-37).

## Acknowledgments

We thank Bailong Xiao (Tsinghua University) for the generous gift of the 9K mutant. This work was supported by NIH grant GM130834 to J.J.L. and Y.L.L.

## Declaration of interests

The authors have no conflict of interests.

## Authors Contributions

Project conception: Y.L.L. and J.J.L. Data acquisition and analyses: Y.L.L., T.D.W., A.B., W.J. and J.J.L. Manuscript writing: J.J.L. with input from all authors.

## Notes

### Competing Interest Statement

The authors have declared no competing interest.

